# Mechano-activation of synovial fibroblasts and macrophages during OA progression in the dynamically stiffening synovial microenvironment

**DOI:** 10.64898/2026.02.16.706240

**Authors:** Sung Yeon Kim, Easton C. Farrell, Kevin G. Burt, Bryan Kwok, Qiushi Liang, Alexander J. Knights, Kate L. Sharp, Vu Nguyen, Lance A. Murphy, Baofeng Hu, Alexandra Kahn, Ling Qin, Lin Han, Tristan Maerz, Robert L. Mauck, Carla R. Scanzello

## Abstract

**Objective:** During osteoarthritis (OA) progression, the synovial membrane undergoes profound structural and compositional remodeling and fibrosis. We sought to elucidate how evolving synovial microenvironmental mechanics during fibrotic remodeling influence cell behavior and drive the progression of synovial pathology.

**Methods:** Skeletally-mature male C57BL/6J mice were subjected to destabilization of the medial meniscus (DMM). To control for surgical confounders, both sham-operated and unoperated mice were included, with evaluation at 4– and 8-weeks. Synovial micromechanics were quantified via atomic force microscopy (AFM). Single-cell RNA sequencing (scRNA-seq), RNA fluorescence *in situ* hybridization (FISH), and flow cytometry were employed to investigate cellular heterogeneity, spatial organization, and crosstalk within fibrotic and non-fibrotic synovial niches.

**Results:** Progressive fibrotic remodeling and marked matrix stiffening were observed in DMM-operated synovium but absent in sham– and un-operated controls. While both sham and DMM joints mounted an acute stromal and immune response to surgery, these changes resolved over time in sham conditions but persisted in DMM synovium. During disease progression, distinct functional subsets of synovial fibroblasts and immune cells emerged, with mechanosignalling pathways and distinct immune cell-fibroblast crosstalk robustly activated within DMM-induced fibrotic microenvironments.

**Conclusion:** This study demonstrates the complex cellular dynamics and crosstalk that differentiate the evolution of the pathological synovial response in the fibrotic DMM condition relative to surgical sham controls. Our findings highlight mechanotransduction as a central mechanism driving OA synovial pathogenesis and underscore the utility of the DMM model as a platform to dissect the molecular underpinnings of synovial fibrosis.

## Introduction

Tissue repair following injury is a complex response, and when injuries are repetitive or chronic, endogenous wound healing responses become dysregulated, leading to fibrosis — a condition marked by excessive deposition of extracellular matrix (ECM). In osteoarthritis (OA), fibrotic remodeling of the joint-lining synovium is a universal pathologic feature that contributes to disease progression, debilitating pain, and joint stiffness [^1–3^].

Fibroblasts and macrophages, two primary cellular components of the synovial membrane, are key mediators of tissue regeneration. The interplay between these two cell types, which can determine whether an injury heals successfully or leads to pathological tissue remodeling, remains poorly understood [^4,5^]. Synovial fibroblasts (SFs) become activated during OA and take on a myofibroblastic phenotype to produce abundant ECM [^6–8^]. These SFs reside near macrophages — a highly heterogeneous synovial cell population whose diversity arises from their polarization, presence of pathology, and ontogeny [^9–11^]. Specifically, tissue-resident macrophages of the intima are derived from an embryonic, monocyte-independent lineage and form a tight junction-mediated barrier that physically restricts inflammatory infiltration. In contrast, both resident and peripherally-derived macrophages occupy the subintimal layer [^9^]. However, it is not yet known how distinct macrophage populations (resident versus recruited) contribute to synovial fibrosis, as well as how crosstalk with SFs changes as they undergo their own fibrotic transition. Addressing these gaps may enhance efforts to identify specific macrophage populations and immune mechanisms that can be targeted for precision OA therapy.

In addition to complex intercellular crosstalk, cells maintain a continuous crosstalk with their physical microenvironment. constantly probing their biophysical milieu through contractile cytoskeletal machinery. Mounting evidence suggests that mechanosensitive pathways modulate the activity of transcription factors (e.g., YAP/TAZ, SRF) to transduce these microenvironmental cues to the nucleus to modulate transcriptional programs [^12,13^]. In joint injury, the synovial membrane undergoes dramatic structural and compositional changes. Namely, increased matrix deposition, disruption of degradative processes, and abnormal production of ECM constituents are thought to stiffen the synovium [^14–16^]. However, it is currently unknown how SFs and macrophages are regulated in the context of such a dynamic biophysical milieu, and whether evolving cell-matrix interactions contribute to aberrant tissue remodeling. While it is well-established that altered matrix mechanics can promote myofibroblast emergence, recent evidence suggests that macrophages are likewise mechanosensitive cells that tune their phenotype in response to biophysical cues, including matrix rigidity, topography, and spatial confinement [^17–20^]. While these studies reveal new perspectives for the design of immunomodulatory biomaterials, the functional consequences of altered microenvironmental mechanics on macrophage phenotype and behavior specifically in the OA synovium, are largely unexplored.

In this study, we employed a murine model of experimental OA that recapitulates chronic joint inflammation and fibrosis to interrogate complex immune-stromal crosstalk and dissect cell-matrix dialogue. Importantly, to address the confounding effects of surgery on synovial inflammation, we incorporated both sham –operated and unoperated controls. We utilized atomic force microscopy (AFM) to characterize synovial micromechanics and single cell RNA sequencing (scRNA-seq), RNA fluorescence *in situ* hybridization (RNA FISH), and flow cytometry to identify and localize discrete immune and stromal cell types in the healthy and OA synovium. This analysis identified potential mechanobiological and paracrine mechanisms responsible for the emergence of pro-fibrotic phenotypes.

## Methods

### Mice

C57BL/6J mice (Jackson Laboratory) were group-housed (4-5 animals/cage) in standard ventilated cages under a 12:12 hour light:dark cycle. Destabilization of the medial meniscus (DMM) surgery was performed on the right knee of 12 week-old male mice while contralateral knees were subjected to sham surgery (medial arthrotomy), as previously described [^21,22^]. Separate age-matched mice served as unoperated controls. Endpoints were defined as 4– or 8– weeks post-surgery. All protocols were conducted in accordance with IACUC protocols at the University of Pennsylvania and Corporal Michael J. Crescenz VA Medical Center.

### Atomic Force Microscopy (AFM)

AFM-nanoindentation was performed on fresh, unfixed 40-μm sagittal cryosections of murine knees (n=10/group) acquired via the Kawamoto’s film-stabilization method [^23^]. Nanoindentation was conducted using polystyrene microspherical tips (*R* ≈ 12.45 μm, nominal *k* ≈ 0.6 N/m, HQ:NSC36/tipless/Cr-Au, cantilever C, NanoAndMore) using a Dimension Icon AFM (Bruker Nano) in 1X PBS with protease inhibitors. Indentation was applied up to a maximum load of ≈120 nN at a 10 μm/s indentation rate. For each animal, indentations were performed on 15-20 distinct locations in synovium to account for spatial heterogeneity of synovium during fibrosis. Effective indentation modulus (*E_ind_*) was computed using the finite thickness-corrected Hertz model, assuming a Poisson’s ratio (ν) of 0.4 for the synovial membrane (**Fig 1D**) [^24,25^].

**Fig 1.**
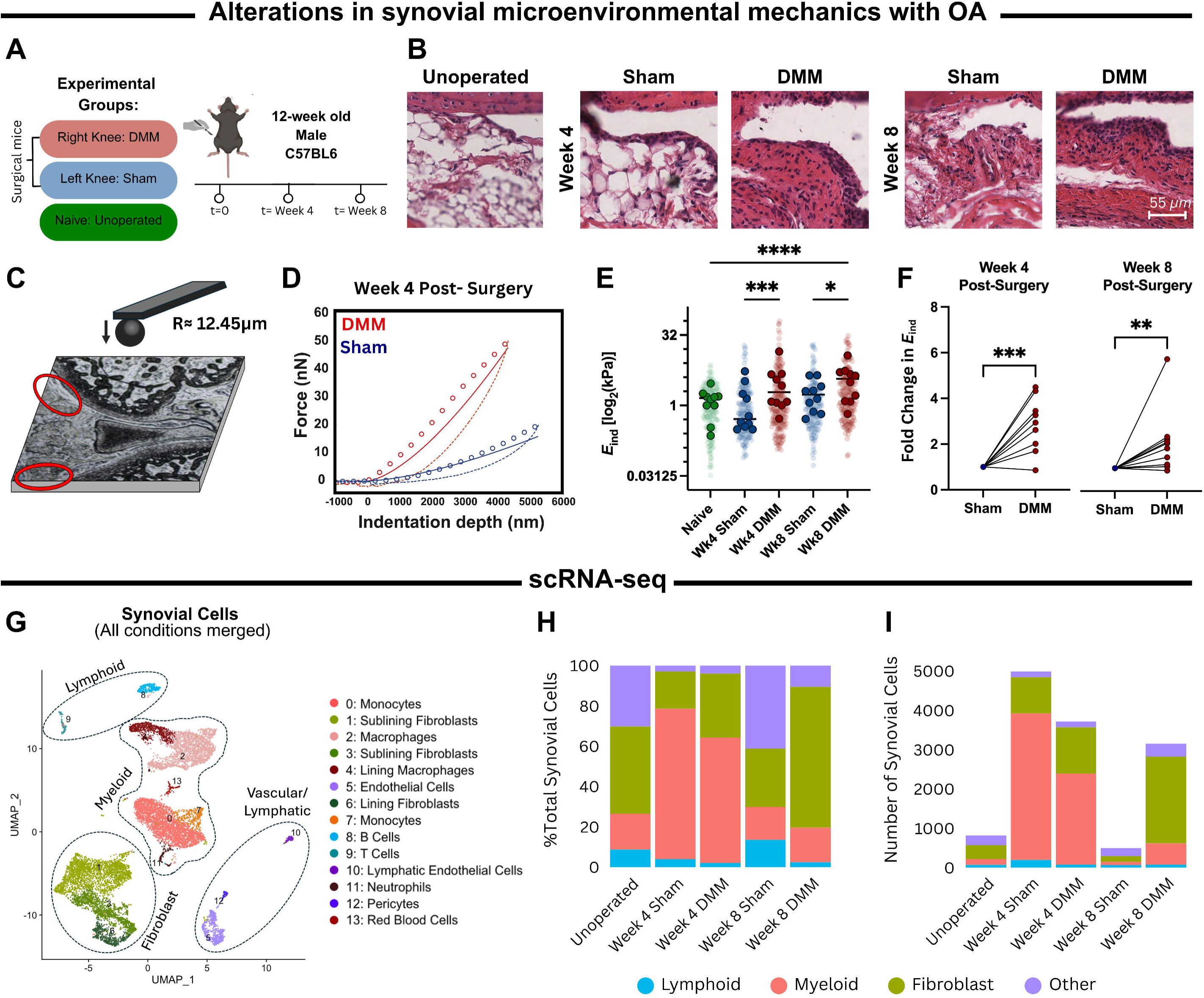
Altered microenvironmental mechanics and cellular landscape in OA synovium. (A) Schematic of experimental design. (B) Representative hematoxylin and eosin (H&E) images of the anterior femoral synovial membrane. (C) Atomic force microscopy (AFM) nanomechanical mapping on cryo-sections of murine knee synovium (*n*= 10 mice per group, 157-202 total indentations per group). (D) Representative indentation force versus depth (*F-D*) curves obtained on Sham and DMM synovia 4 weeks post-surgery (measured in PBS with protease inhibitors, 10µm/s rate). Red and blue solid lines depict finite thickness-corrected Hertz model fitted to the entire loading portion (dotted line) of the DMM and Sham *F-D* curves, respectively. Dashed lines represent the unloading portion of *F-D* curves. (E) Superplot of the micromoduli of the synovial membrane (n=10/condition) (F) Fold change in average synovial micromoduli between Sham and DMM joints for each timepoint (G) UMAP projection of all synovial cells from Sham-, DMM-, and un-operated knees (13,177 cells, n=15 biological replicates per condition) (H) Proportional breakdown and (I) total abundance of major synovial cell types in each condition.

### Synovial Tissue Harvest

To isolate cells for flow cytometry and scRNA-seq, synovial tissue was harvested from the medial, lateral, and anterior compartments of the knee joints, inclusive of the fat pad, and digested for 30 minutes in RPMI with Liberase (1 U/mL) and DNase I (200 μg/mL) at 37°C, with intermittent agitation. Cell suspensions were then filtered through 40-μm strainers and washed twice with RPMI containing 10% fetal bovine serum. For scRNA-seq, synovial cells were resuspended in a sterile-filtered BSA solution (1% w/v), while for flow cytometry synovial cells were reconstituted in commercial FACS buffer (BioLegend).

### Single-cell RNA Sequencing (scRNA-seq)

Knee synovial cell suspensions from 15 individual mice were combined for each experimental condition (unoperated, sham and DMM 4 weeks, sham and DMM 8 weeks). Two replicates (i.e., discrete sequencing experiments) were performed for each condition, yielding a total of ten samples. Single cell suspensions (>90% viable) were immediately submitted to the Children’s Hospital of Philadelphia (CHOP) Center for Applied Genomics for processing through the 10x Genomics pipeline (Chromium Next GEM Single Cell 3’ Kit v3.1), according to manufacturer instructions. Libraries were uniquely indexed using the Chromium Dual Index Kit, pooled, and subsequently sequenced on the Illumina NovaSeq 6000 platform in a paired-end, dual indexing run, generating ∼20,000 mean reads per cell. Data were then processed using the CellRanger pipeline (10x Genomics, v.6.1.2) for demultiplexing, read alignment to the mm10 (*Mus musculus*) transcriptome, and creation of feature-barcode matrices.

<u>Histology:</u> Hind limbs from mice were harvested and fixed for 48 hours in 10% neutral-buffered formalin, decalcified in 10% EDTA, and paraffin processed. Sagittal sections (10 µm) spanning the medial compartment were sectioned and stained with Hematoxylin and Eosin (H&E), then scanned on an Aperio ImageScope (Leica). Semi-quantitative synovitis grading was performed at 20x magnification by three blinded observers according to our established scoring method [^26^].

### Flow Cytometry

After enzymatic tissue digestion, single-cell suspensions were washed with commercial FACS buffer and nonspecific binding was blocked by incubation with 2 µl of TruStain FcX (BioLegend) for 20 minutes. Cells were stained with fluorescently conjugated antibodies (**Supplementary Table 1**) for 30 minutes on ice in the dark, washed, and resuspended. Unstained controls and FMO controls were included to define negative and positive gating boundaries for markers. SSC beads (BioLegend) were used to set voltages and compensation. Flow cytometry was performed on a LSRFortessa machine using FACSDiva software for data acquisition, and data were analyzed with FlowJo v10.9.0 (BD/Treestar).

### Tissue Sectioning and Hybridization Chain Reaction (HCR) RNA Fluorescence *In Situ* Hybridization (FISH)

Harvested hindlimbs were fixed in RNase-free 4% paraformaldehyde solution for 24 hours at 4°C, then washed and decalcified in freshly prepared Morse’s solution (10% sodium citrate, 22.5% formic acid) at room temperature for 72 hours. Knee joints were embedded in paraffin and sectioned at 7-µm thickness. *In situ* hybridization was performed using the HCR^TM^ RNA-FISH assay by Molecular Instruments, according to manufacturer’s instructions. Custom *Adgre1*-B1, *Trem2*-B2, and *Cx3cr1*-B3 probes (Molecular Instruments) were used to visualize the expression of F4/80 (NM_010130.5), *Trem2* (NM_031254.4), and *Cx3cr1* (NM_009987.4) respectively. Stained sections were imaged on the Zeiss Axio Scan Z1.

## Results

### The microenvironmental mechanics of synovium change with OA progression

We first sought to assess structural and micromechanical dynamics of OA synovium during disease progression. We employed the murine DMM model (**Fig 1A**), which reproducibly induces key pathologic features (e.g. cartilage degradation, synovial inflammation, and fibrosis) and resembles slowly progressing human OA [^21^]. H&E staining corroborated the progression of synovial fibrosis in DMM-operated mice at 4– and 8– weeks post-surgery (**Fig 1B**), along with synovial lining hyperplasia and increased sub-intimal cellularity (**Supplementary Fig 1A**). To correlate histological changes with microenvironmental mechanics, we performed AFM-nanoindentation on the subintimal layer of the anterior synovial membrane of DMM-, sham-, and unoperated knee joints (**Fig 1C**). At both 4-weeks and 8-weeks post-surgery, the synovium in DMM-operated joints showed a significant increase in moduli (*E_ind_*) (1.5 ± 0.4 kPa versus 3.6 ± 1.1 kPa for Week 4– sham and DMM, respectively; 2.1 ± 0.5 kPa versus 3.2 ± 0.7 kPa for Week 8– sham and DMM, respectively; mean ± 95% CI) (**Fig 1E**). Notably, this represented an average 2.1– and 2.9– fold increase in DMM joints compared to sham at 4-weeks and 8-weeks post-surgery, respectively (**Fig 1F**), consistent with the degree of matrix stiffening reported in fibrotic lung disease [^27^]. DMM synovium was also consistently stiffer than unoperated synovium (1.3 ± 0.2 kPa). These results demonstrate fibrotic remodeling in the OA synovium is evident through both histology and alterations in microenvironmental tissue mechanics.

### Cellular expansion and diversification accompany synovial remodeling during OA progression

To dissect the dynamic cellular landscape within the remodeling biophysical milieu, we next performed scRNA-seq of synovial cells isolated from DMM, sham, and unoperated mice (**Fig 1G**). SFs, myeloid cells, and endothelial cells constituted the majority of cells within the healthy synovium, consistent with previous reports [^6,8^]. At the 4-week timepoint, the myeloid population dramatically expanded in both sham and DMM knees, in response to surgical insult. By 8 weeks, this expansion subsided in both groups, though only the sham synovium returned to its baseline cellular composition, while abundance of SFs and macrophages in DMM synovium remained higher compared to unoperated controls (**Fig 1H & 1I**). These data show that joint injury results in dynamic changes in stromal and immune cell populations that ultimately resolve towards baseline in sham but persist in DMM joints.

### Functionally distinct SFs subsets emerge and persist in OA synovium

To understand synovial stromal changes with disease progression, we evaluated global changes in SF transcriptional activity using pseudobulk expression profiles for each condition. Multidimensional scaling (MDS) visualized these profiles and demonstrated that at 4-weeks post-surgery, SFs from both surgical groups diverged markedly from unoperated controls. By 8-weeks, sham SFs shifted back toward the transcriptional state of unoperated joints, while DMM SFs remained distinctly separated, indicating persistent aberrant transcriptional activity (**Fig 2A**). Consistent with this, many differentially expressed genes (DEGs; p_adj_<0.05) were identified at 4-weeks compared with unoperated controls (977 and 1,077 DEGs in sham and DMM joints, respectively). This divergence attenuated by 8 weeks, particularly in sham joints (67 and 186 DEGs in sham and DMM joints, respectively) (**Fig 2B**). Hierarchical clustering and Pearson correlation analysis further reflected these temporal patterns (**Fig 2C**). Comparison of DEGs between sham and DMM SFs at both times, along with the expression levels of those genes in unoperated SFs, revealed a strong similarity between unoperated and sham groups, with notable deviations in the DMM group (**Fig 2D**). Both surgical groups showed peak mechano-activation (**Fig 2E**) and matrix turnover activity (**Fig 2F**) at 4-weeks. Collagen genes previously linked to OA (types I, III, VI) were upregulated in both surgical groups [^28,29^]. However, sham SFs, exhibited a balance between matrix deposition and degradative enzyme gene expression, with elevated nascent matrix genes (e.g. *Col3a1, Fn1, Has1*) accompanied by higher expression of matrix metalloproteinases (MMPs) and lower MMP inhibitors (e.g. *Timp1*). In contrast, DMM SFs skewed towards net matrix accumulation.

**Fig 2.**
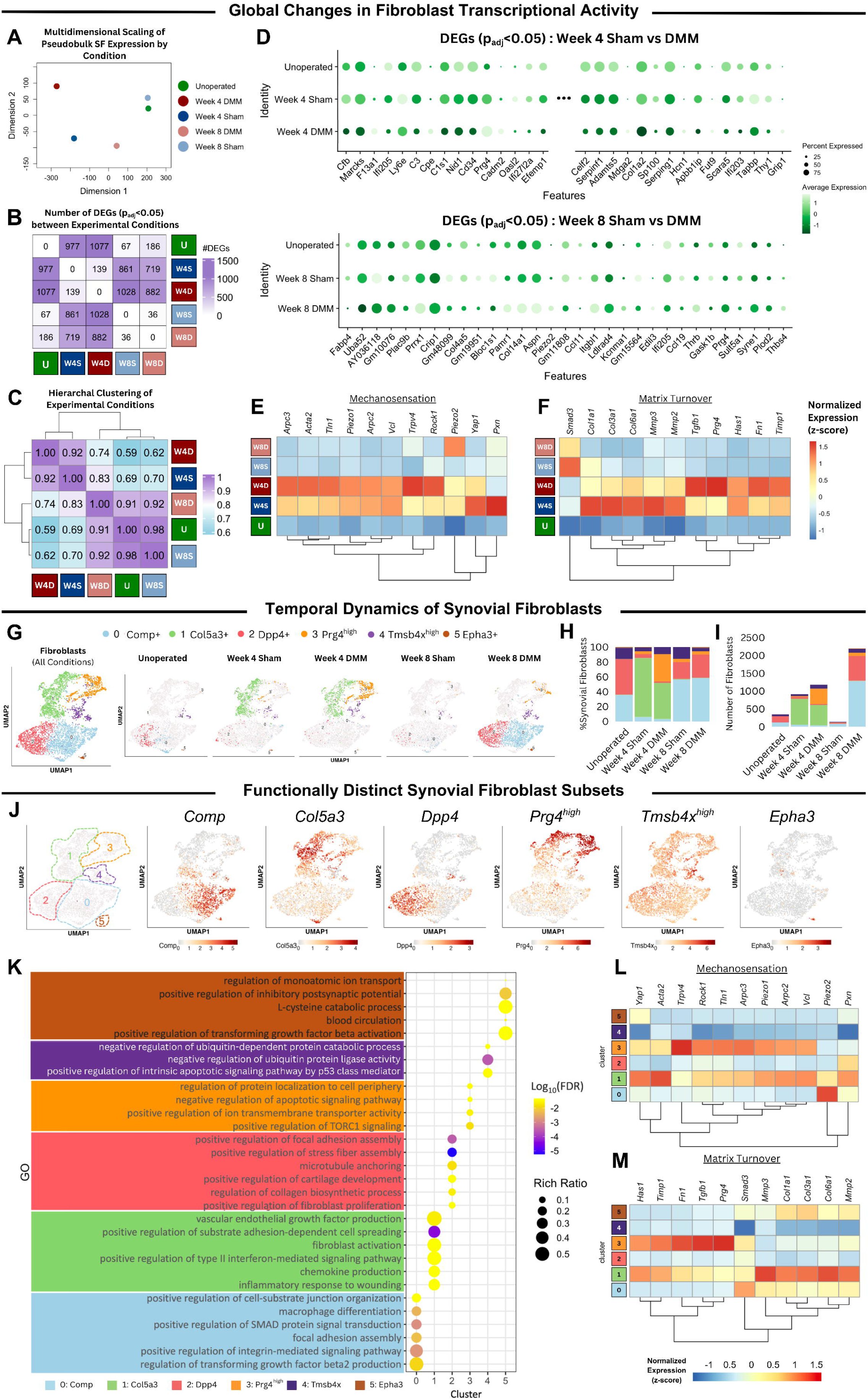
Identification of functionally distinct synovial fibroblast subtypes. (A) Multidimensional scaling (MDS) of pseudobulk fibroblast expression by experimental conditions. (B) Heatmap showing the number of differentially expressed genes (DEGs; p_adj_<0.05) between experimental conditions [U= Unoperated, W4S= Week-4 Sham, W4D= Week-4 DMM, W8S= Week-8 Sham, W8D= Week-8 DMM]. (C) Heatmap displaying the correlation of gene expression for all pairwise combinations of experimental conditions in the dataset. (D) Bubble plot showing the top DEGs (p_adj_ <0.05) between Sham and DMM SFs at 4– (top) and 8-weeks (bottom) post-surgery. Expression levels for the same genes in unoperated SFs are also shown. Heatmaps showing expression (z-scores) of (E) mechanosensitive and (F) matrix remodeling genes across experimental conditions. (G) UMAP projection showing the temporal evolution of synovial fibroblasts isolated across experimental conditions (*n*=15 biological replicates per condition). (H) The proportional breakdown and (I) total number of SF subtypes in each condition. (J) Feature plots of marker genes used to delineate each SF subpopulation. (K) Pathway analysis using Gene Ontology (GO) Biological Processes annotating unique functional terms for each cluster. Heatmap of expression levels (z-score) of (L) mechanosensitive and (M) matrix remodeling genes by SF subtype. (N) Box plots showing the ratio of apoptotic to cell cycle module scores for each SF subcluster.

Previous work demonstrated functionally-distinct SF subsets in noninvasive joint trauma [^6^]. Here, we extend these findings by characterizing SFs during resolving and non-resolving (OA-developing) surgical trauma. To enable more granular interrogation, we computationally subset and re-clustered SFs and related mesenchymal cells, revealing six sub-clusters defined by canonical marker-gene expression (**Fig 2G & 2J**). As expected, SF markers *S100a4, Pdpn, Vim,* and *Pdgfra* were broadly expressed, with a distinction between *Thy1*+ subintimal (*Comp*+, *Col5a3*+, *Dpp4*+, and *Epha3*+) and *Thy1*-intimal SFs (*Prg4^high^* and *Tmsb4x^high^*) (**Supplementary Fig 1D, E**) [^30,31^]. Strikingly, while *Comp*+ and *Dpp4*+ SFs dominated healthy synovium, they declined at 4-weeks post-surgery, while a *Col5a3*+ SF population emerged in both surgical groups. Consistent with prior descriptions of myofibroblast-like SFs, *Col5a3*+ cells also expressed *Acta2* (21.3%), *Tagln* (17.9%), and *Lrrc15* (44.4%) (**Supplementary Fig 2I)**, implying a myofibroblast-like phenotype [^6, 32^]. Concurrently, only DMM synovium displayed increased *Prg4^high^* lining SFs, aligning with chronic lining hyperplasia in OA [^1^]. By 8-weeks, the transient *Col5a3*+ population dwindled in both sham and DMM, with sham returning to near baseline. Although DMM also returned towards baseline proportions, total SF numbers were increased 6.4-fold, including sustained elevations in *Prg4* ^high^ SFs (**Fig 2H & 2I**). These SF changes reflect hallmark pathologic features of OA synovium –lining hyperplasia and subintimal cellularity—with the increase in *Prg4^high^* lining SFs mirroring findings in the non-invasive ACL rupture model [^6^]. Trajectory analysis (Monocle3). identified *Dpp4*+ progenitors as precursors to transient *Col5a3*+ and *Prg4* ^high^ SFs (**Supplementary Fig 1G**). Interestingly, the transient *Col5a3*+ SFs showed no significant apoptotic gene signatures, suggesting that their disappearance at 8-weeks may reflect further phenotypic differentiation rather than cell death (**Supplementary Fig 1H**).

Having characterized the temporal dynamics of SF subsets, we next examined whether the unique transcriptomic profiles of each cluster reflected distinct functional roles. Gene Ontology analysis identified unique functional terms for each cluster (**Fig 2K**). Notably, both the transient *Col5a3*+ and *Prg4^high^* clusters demonstrated elevated expression of genes encoding mechanosensitive proteins and transcription factors relative to other SF subsets (**Fig 2L**). Despite this shared feature, the transient *Col5a3*+ subset appeared to maintain functional balance, serving as both a major producer of collagenous matrix components and a key source of MMPs. *Prg4^high^* SFs were the predominant source of *Tgfb1,* suggesting a potential role as a paracrine pro-fibrotic driver (**Fig 2M**). Analysis of GO pathways related to mechanotransduction (GO:2001046-positive regulation of integrin-mediated signaling pathway) and matrix turnover (GO:0032965-regulation of collagen biosynthetic process) further supported that transient *Col5a3*+ and *Prg4^high^* SFs are most engaged in mechanosensitization and matrix remodeling (**Supplementary Fig 1F**). These pathways also remain significantly elevated in *Dpp4*+ and *Comp*+ subsets, which may underscore the significance of their proliferative expansion within the fibrotic synovium.

Collectively, these findings highlight the functional diversity of SF subsets, with distinct transcriptional patterns that suggest specialized roles. Comparisons between sham and DMM highlight both shared and divergent features that likely reflect context-dependent functions related to disease-resolution or progression. Importantly, the identification of enriched, subsets—*Col5a3*+ and *Prg4^high^* SFs—with aberrant, highly mechano-active transcriptional patterns implicate overactivation of SF mechanotransduction as a potential driver of synovial pathology in disease.

### A divergent immune landscape emerges in the OA synovium

Having characterized the disease-associated diversity of the stromal population, we next evaluated the immune cell landscape in healthy and injured synovium. We computationally subset and re-clustered immune cells (*Ptprc*+, *Pdgfra*-), revealing diverse innate and adaptive populations, including monocytes, macrophages, T and B cells, dendritic cells, and neutrophils (**Fig 3A, 3B**). Immune cell abundance increased in both sham and DMM at 4-weeks, with monocytes and macrophages being the predominant constituents numerically and proportionally (**Fig 3C**). While sham joints resolved towards baseline by 8-weeks, the DMM synovium retained the expanded immune cell population (∼185% increase above unoperated). At this timepoint, DMM showed both numerical and proportional enrichment of *Trem2*+ *Cx3cr1*+ tissue-resident macrophages (**Fig 3D**). Multi-color flow cytometry using CD45, F4/80, TREM2, and CX3CR1 confirmed the increased representation of this subset in the 8-week DMM group (**Fig 3J**). RNA FISH analysis then localized this *Trem2*+*Cx3cr1*+ macrophage expansion to the synovial intima (**Fig 3I, Supplementary Fig 2**).

**Fig 3.**
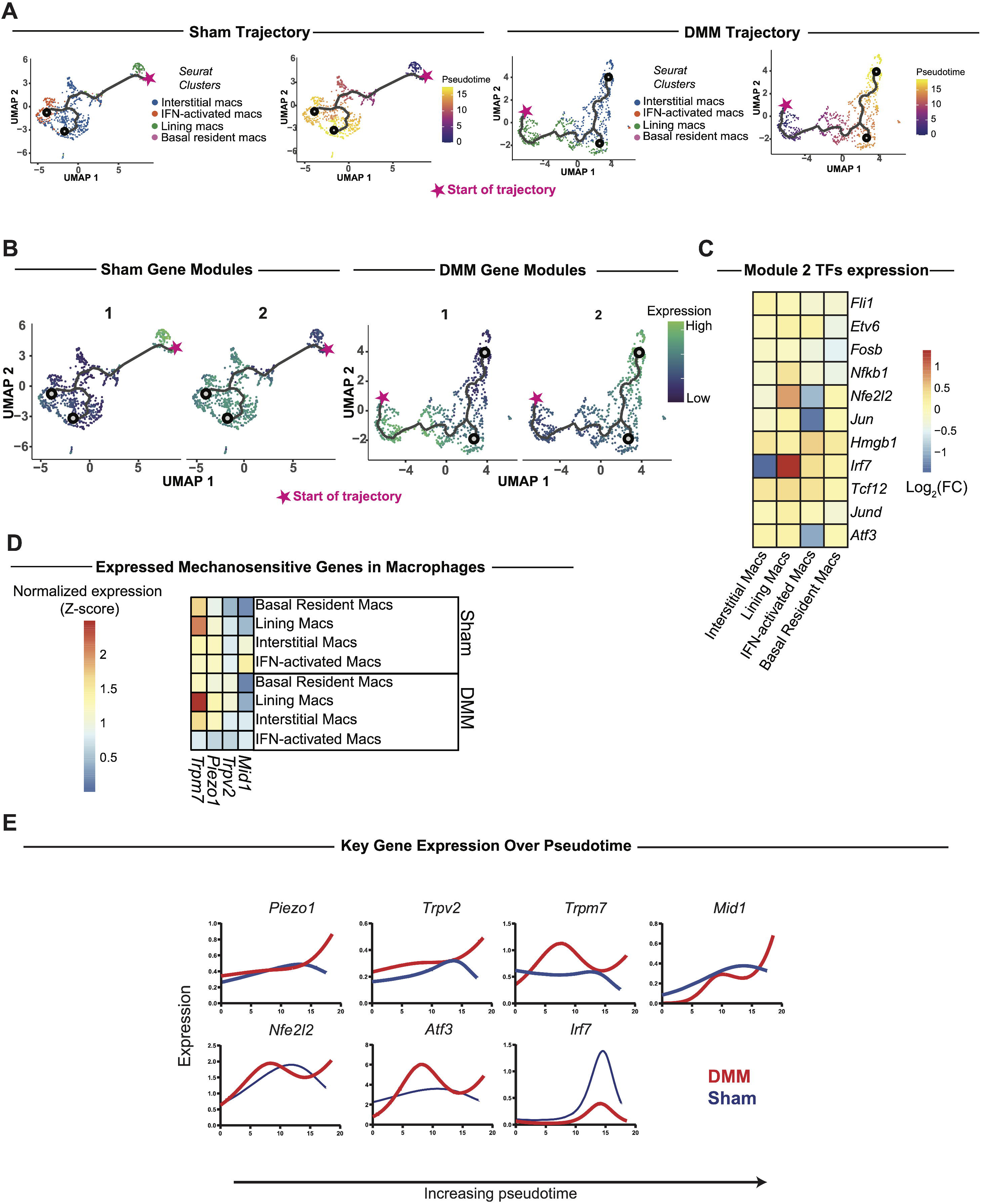
The dynamic immune landscape in the osteoarthritic synovium. (A) UMAP projection illustrating the temporal evolution of CD45+ synovial cells subsets. (B) Violin plots showing expression levels of immune cell marker genes across distinct clusters. (C) Proportional breakdown and (D) total cell counts of CD45+ subtypes in each experimental condition. Violin plots showing Pro-Inflammatory (E, G) and Pro-Fibrotic (F, H) module scores across immune cell subtypes or experimental conditions. (I) Representative RNA FISH images localizing lining macrophages; dashed lines demarcate the synovial tissue and arrows indicate the synovial intima. (J) Flow cytometry quantification of the proportion of TREM2+ CX3CR1+ lining macrophages out of all CD45+ cells as detected. Exhaustive list of significantly (p<0.05) differentially expressed genes in *Trem2+ Cx3cr1+* lining macrophages at (K) 4-weeks and (L) 8-weeks post-surgery Sham and DMM synovium.

To infer functional roles of each immune cluster, we computed pro-inflammatory module scores using genes involved in myeloid M1-like and lymphoid Th1/Th17-like responses (**Fig 3E**), and pro-fibrotic module scores using genes involved in inducing fibrotic phenotypes, including *Tgfb1, Pdgfb, Osm, Vegfa, Mmp9,* and *Csf1r* (**Fig 3F**). Classical intermediate monocytes, non-classical monocytes, and IFN-activated macrophages had the highest pro-inflammatory scores. Both pro-inflammatory and pro-fibrotic signatures were most enriched at 4-weeks in sham and DMM (**Fig 3G, 3H**). By 8 weeks, however, Wilcoxon tests with Benjamini-Hochberg correction revealed a significant increase in the pro-fibrotic module score in DMM synovium compared to sham controls (*p_adj_*=2.24×10^-7^). Although global comparisons of immune cell composition and pro-fibrotic/inflammatory activity revealed broad similarities between sham and DMM joints — especially at 4-weeks — granular transcriptomic analysis uncovered divergent patterns. *Trem2*+ *Cx3cr1*+ macrophages exhibited the most DEGs between sham and DMM at 4-weeks (n=27; **Fig 3K**), compared to just 11 at 8-weeks (**Fig 3L**). In DMM, these macrophages upregulated genes involved in ECM remodeling (*Col3a1, Fn1, Spp1*), lysosomal degradation and protease regulation (*Ctsd, Lgals3, Ctla2a*), and innate immune activation (*Clec4d, Ms4a7, Cd84, Fcrls, Tlr7*). Collectively, these findings suggest *Trem2*+*Cx3cr1*+ macrophages orchestrate a transcriptional program promoting fibrosis and inflammation in the DMM synovium.

### Distinct macrophage differentiation trajectories define DMM versus sham synovium

Having identified the plasticity of macrophage abundance and phenotypes in sham and DMM synovium, we next subset macrophages computationally and analyzed their differentiation trajectories using Monocle3. Sham and DMM exhibited consistent differentiation trajectories from basal resident (defined as *Trem2*+ *Cx3cr1*+ macrophages in unoperated samples) to interstitial and IFN-activated macrophages, via lining macrophage intermediates (**Fig 4A**). We identified two modules of genes with shared expression patterns over pseudotime. Module 1 contained genes enhanced in basal and lining macrophages (low pseudotime) and Module 2 contained genes enhanced in interstitial and IFN-activated macrophages (high pseudotime) (**Fig 4B**).

**Fig 4.**
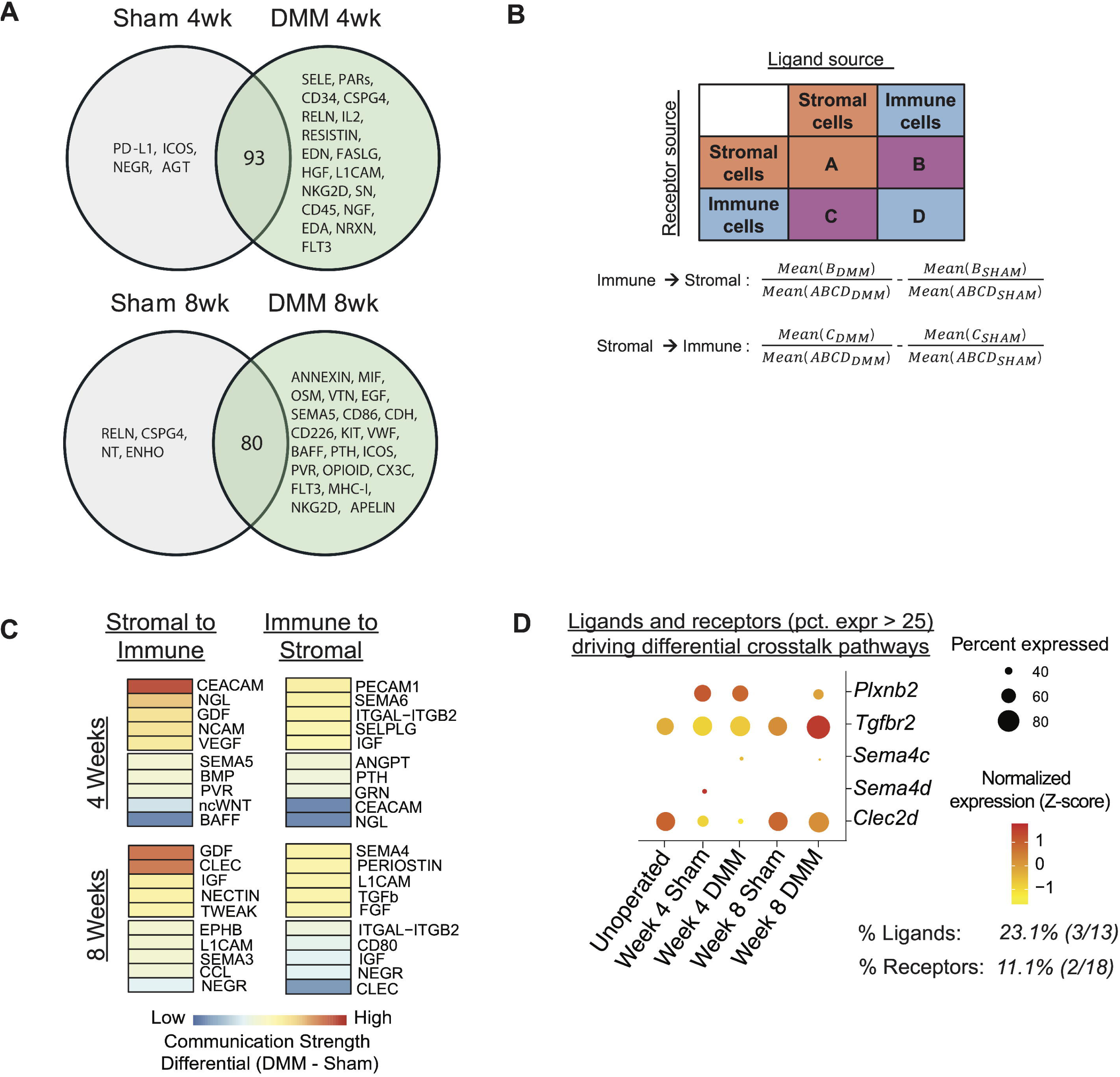
Macrophage differentiation trajectories in OA synovium. (A) Monocle3-inferred trajectories from sham and DMM basal resident macrophages to interstitial and IFN-activated macrophages and corresponding and pseudotime feature plots. (B) Feature plots of sham and DMM gene modules that similarly vary along pseudotime. (C) Relative expression of significant, direct-annotation derived transcription factors from RcisTarget (NES > 3.0) (D) Transcript expression of mechanosensitive genes substantially expressed in either sham or DMM macrophages (at least one cluster with average expression > 0.5 counts). (E) Pseudotime regression plots of sham and DMM macrophage expression of key mechanosensitive genes and significant transcription factors.

To identify potential molecular regulators driving differentiation trajectories, genes from Module 2 from each condition were analyzed with RcisTarget, which identifies transcription factor binding sites shared among module genes. Eleven annotation-derived transcription factors associated with enhanced expression of genes in Module 2 were found in either sham or DMM, with the majority of these also upregulated in DMM compared to sham macrophages (**Fig 4C**). Since macrophage mechanosensation is poorly characterized in OA, we next screened expression of more than 30 genes involved in mechanosensation and identified 4– *Piezo1, Trpm7, Trpv2,* and *Mid1 –*appreciably expressed in synovial macrophages (**Fig 4D, Supplementary Fig 2B**). Pseudotime regression plots of these transcription factors and mechanosensitive genes showed that *Piezo1, Trpv2, Atf3,* and *Nfe2l2* remained highly expressed at the termini of the differentiation trajectories in DMM but were expressed at low levels at the termini in sham (**Fig 4E**), suggesting that expression of these genes may contribute to the differential synovial macrophage activity observed in OA.

### Crosstalk between synovial fibroblasts and immune cell populations

A hallmark of OA is chronic inflammation, thought to be driven by pathological crosstalk between stromal and immune cells. We used CellChat to identify cell-cell communication axes differentially active in sham and DMM [^33^]. At both 4– and 8-weeks, we identified significant crosstalk pathways exclusive to sham or DMM (**Fig 5A**). In sham, PD-L1 (programmed death ligand-1), ICOS (inducible T-cell co-stimulator), NEGR (neuronal growth regulator), and AGT (angiotensinogen) signaling were exclusively identified at 4 weeks, and RELN (reelin), CSPG4 (chondroitin sulfate proteoglycan-4), NT (neurotensin), and ENHO (energy homeostasis associated) were exclusively identified at 8 weeks. DMM exhibited several exclusive and significant crosstalk pathways, as well, including VWF (Van Willebrandt Factor), KIT (receptor tyrosine kinase), SEMA5 (semaphorin 5A), and NGF (nerve growth factor). These pathways mediate endothelial development, immune cell activation, axonal guidance, pain, and nerve growth, which are established pathologies in OA.

**Fig 5.**
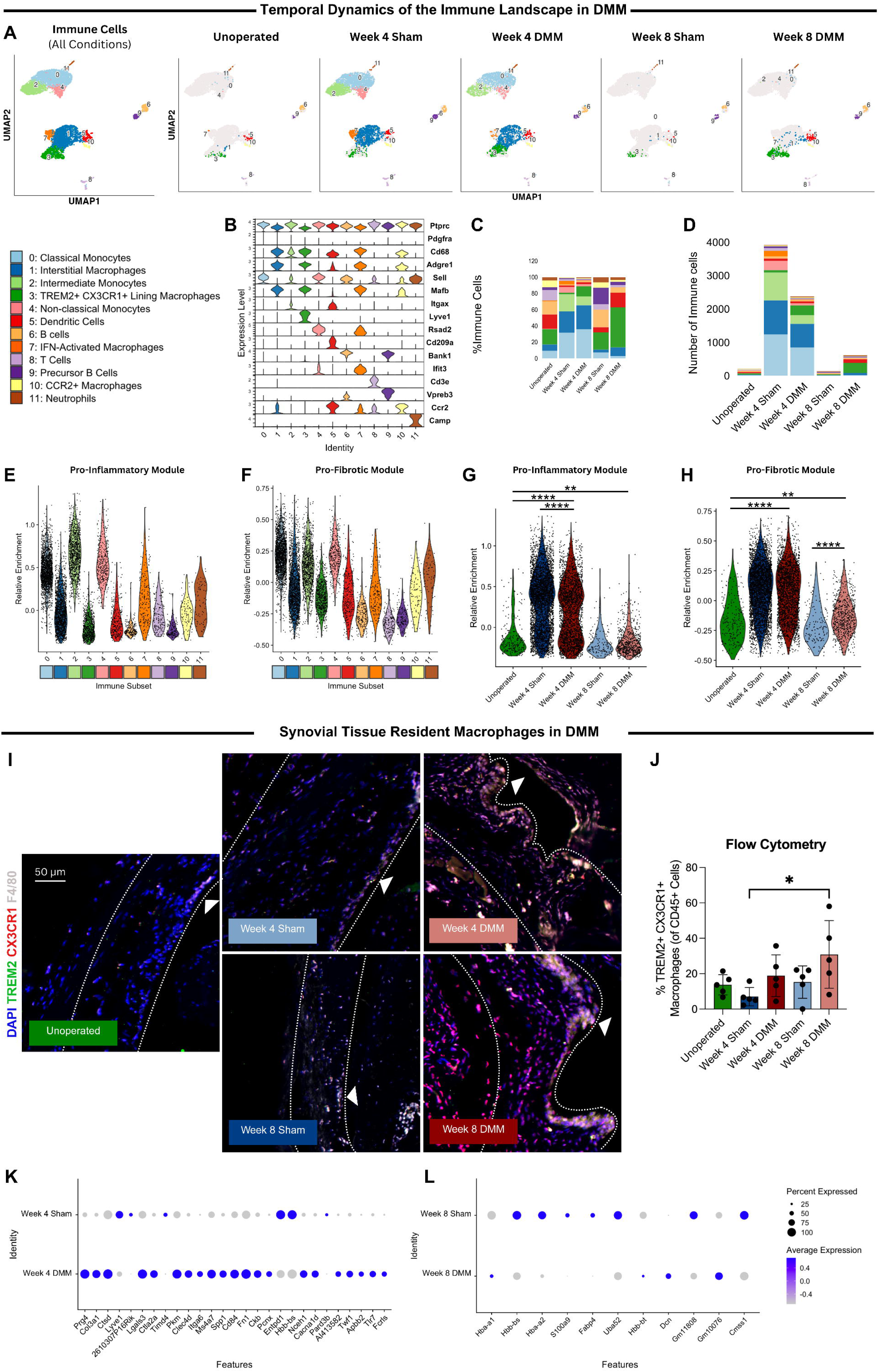
Differential crosstalk in DMM and sham Synovium. (A) All unique significant CellChat pathways for sham and DMM stromal and immune cell communications (p < 0.05). (B) Formulas for calculation of differential crosstalk communication strength. (C) Top 5 highest (enhanced in DMM) and top 5 lowest (enhanced in sham) CellChat pathways ranked by differential crosstalk communication strength. (D) Z-score and percent of positively expressing cells of ligands and receptors underpinning the top two up– and down-regulated CellChat pathways per time point (pct > 0.25 in at least one condition).

**Fig 6.**
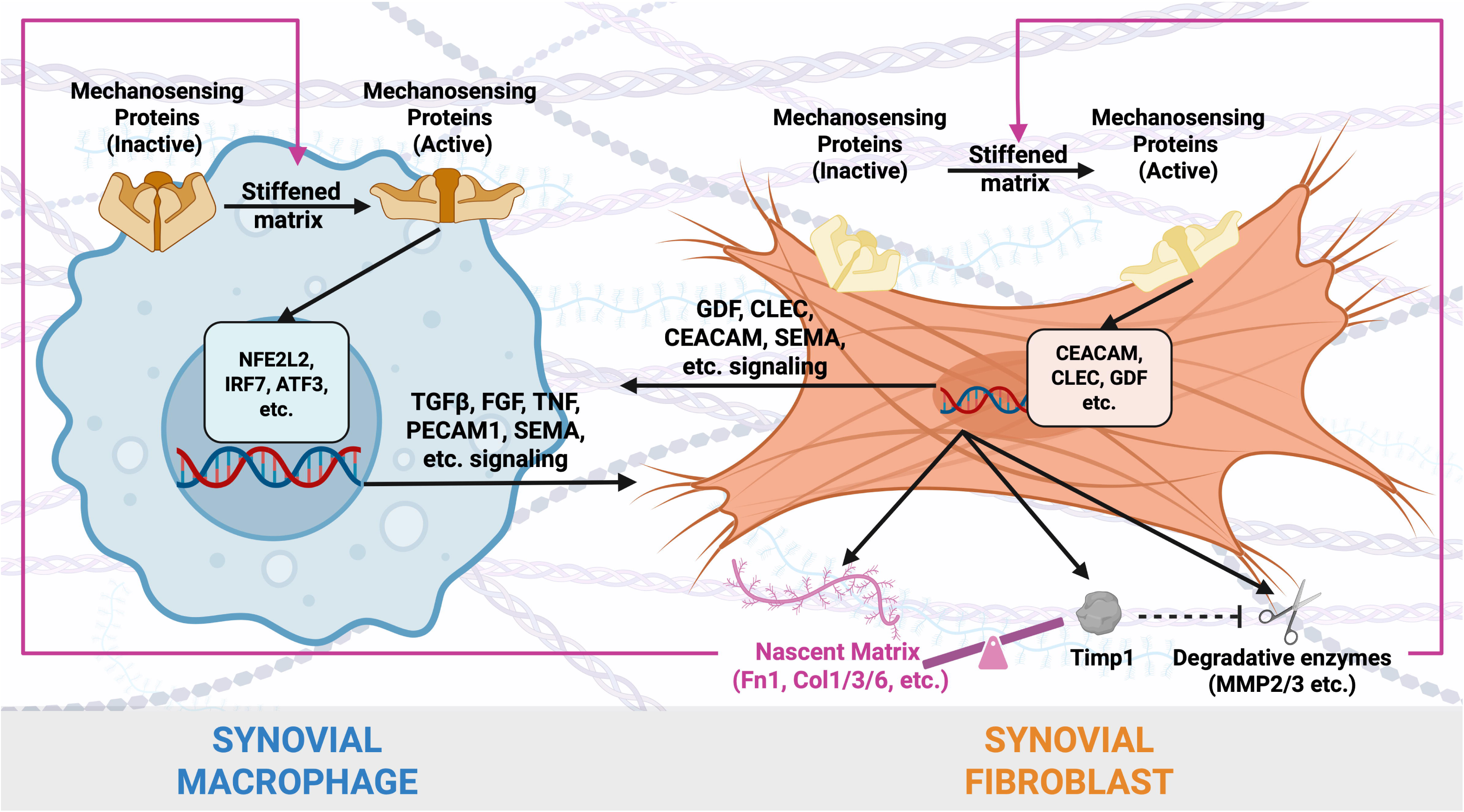
Schematic illustrating proposed mechanotransductive processes involved in synovial fibrogenesis in OA.

To identify crosstalk mechanisms differentially active between sham and DMM, we ranked all significant crosstalk pathways by the proportion of signaling in each direction (stromal-to-immune or immune-to-stromal) and calculated differential crosstalk scores (**Fig 5B**). For both communication directions, several pathways were more strongly engaged in either sham or DMM (**Fig 5C**). GDF (growth differentiation factor), NGL (netrin-g ligand), CLEC (c-type lectin), and CEACAM (carcinoembryonic antigen-related cell adhesion molecule) were most differentially active in stromal-to-immune signaling in DMM while SEMA4 (semaphorin-4), SEMA6 (semaphorin-6), PERIOSTIN (periostin), and PECAM1 (platelet and endothelial cell adhesion molecule-1) were most differentially active in immune-to-stromal communication. BAFF (B cell activating factor), NGL and CEACAM were most differentially active in sham at 4 weeks, whereas CLEC, NEGR and CCL (chemokine ligand) signaling pathways were most active in sham at 8 weeks. Of the ligands and receptors underpinning top crosstalk pathways, only 3 ligands and 2 receptors — *Plxnb2*, *Tgfbr2*, *Sema4c*, *Sema4d*, *Clec2d* — were expressed in at least 25% of cells in any condition, and thus may represent therapeutic targets to assuage the pro-fibrotic response in DMM synovium (**Fig 5D**).

Taken together, our findings demonstrate a distinct pattern of crosstalk between stromal and immune cells in sham and DMM synovia, which could drive resolving wound healing versus pro-fibrotic processes, respectively.

## Discussion

The synovium undergoes profound structural and functional changes following joint injury, yet mechanisms orchestrating these changes are incompletely defined. Here, we comprehensively map SFs and immune cells at single-cell resolution in healthy and injured joints using a widely adopted surgical OA model. Our analyses identified functionally distinct cellular subsets shaped by their microenvironment, implicating mechanotransduction as a central driver of synovial pathogenesis, and position the DMM model as a robust platform in which to dissect the mechanisms of fibrotic disease progression versus wound healing within the synovium.

While recent efforts have described synovial cellular complexity in OA models, the impact of altered tissue mechanics on synoviocyte behavior remains less explored. Biomechanical alterations of the synovium concomitant with fibrosis have long been assumed, but the extent of synovial ECM stiffening has not been rigorously quantified. We previously demonstrated synovial stiffening in a canine PTOA model [^34^]; here, we extend these findings to mice, demonstrating fibrosis-associated ECM mechanical stiffening, with important implications for resident and infiltrating cells within the synovium. Increased matrix stiffness can promote the emergence and survival of contractile myofibroblasts with pathogenic phenotypes, characterized by excessive ECM production and loss of homeostatic SF functions—such as secreting joint lubricants [^7^]. In this study, we identified two highly mechano-responsive SF subsets—*Prg4^high^*and *Col5a3*_— with elevated remodeling and mechanotransductive gene expression. Interestingly, *Prg4^high^*SFs were persistently expanded in DMM (consistent with previous reports [^6,8^]) and transcriptionally biased toward matrix production, suggesting they may represent a promising therapeutic target for mitigating synovial fibrosis in OA.

These two mechano-active SF subsets dominated at 4-week post-surgery but were replaced by a much-expanded complement of the baseline *Comp*+ and *Dpp4*+ SFs by 8-weeks. *Comp*+ SFs also exhibited active mechanosignaling and contributed to collagen biosynthesis. Interestingly, *Comp*+ fibroblast expansion has been observed in synovial tissue biopsies of treatment-refractory rheumatoid arthritis [^35^]. *Comp*+ and *Dpp4*+ were highly expanded within the fibrotic DMM niche, compared to sham, suggesting that biophysical cues regulate their proliferation and survival. Prior work. showed that *Dpp4*+ mesenchymal progenitors reside in the sublining layer and function as a fibroblast “reservoir”, giving rise to other specialized stromal cells in both physiological and pathological contexts. More recently, studies from the Qin and Maerz groups demonstrated that *Dpp4*+ cells serve as precursors for *Prg4*+ lining fibroblasts during postnatal growth and OA progression, but not vice versa [^6,8^]. Our current trajectory analyses revealed that the transient *Col5a3*+ SF may also arise from *Dpp4*+ progenitors, but their subsequent disappearance—coinciding with expansion of the *Dpp4*+ population at later timepoints—is difficult to reconcile. Further *in vivo* and mechanistic studies are warranted to clarify whether *Col5a3*+ subsets exhibit plasticity to revert or de-differentiate, or whether distinct waves of *Dpp4*+ progenitors are mobilized in response to evolving microenvironmental cues [^31,36^]. Deciphering these lineage trajectories and mechanical dependencies will be critical to understanding fibroblast heterogeneity and informing novel precision OA therapeutics.

Integrating single-cell RNA sequencing, AFM, flow cytometry, and RNA FISH, we show that DMM recapitulates sustained fibrosis, while the sham-operated joints fully resolve stromal and immune cell responses after surgical injury. This model offers an ideal system to dissect mechanisms of fibrotic persistence versus resolution [^37^]. By 8 weeks, sham SFs regained homeostatic transcriptional profiles suggestive of a wound-healing trajectory, while DMM SFs retained aberrant transcriptional activity favoring ECM accumulation, highlighting a failure to resolve injury-induced responses. Effective repair (in sham) appears to rely on SFs balancing synthesis and degradation, whereas fibrosis (in DMM) reflects the preponderance of dysregulated, matrix-accumulating SFs.

Our data also identified that immune-stromal interactions diverged in DMM and sham: both conditions elicited acute myeloid inflammatory response that resolved in sham but persisted through 8 weeks in the DMM synovium highlighting chronic immune activation as a hallmark of fibrotic persistence. Particularly striking was the enrichment of *Trem2+Cx3cr1+* tissue-resident macrophages in DMM, localized to the synovial lining. Although these cells have been associated with protective functions in rheumatoid arthritis, their persistent expansion in OA suggests pathogenic adaptation to chronically stiffened and inflammatory microenvironments.

Our trajectory analysis revealed conserved differentiation trajectories from basal to activated states across both conditions. However, pseudotime analysis uncovered condition-specific transcriptional programs, including a role for *Piezo1, Trpv2*, and transcription factors such as *Atf3* and *Nfe2l2* in late-stage macrophages from DMM compared to sham. These findings align with recent reports demonstrating enhanced inflammatory activity and impaired wound healing transcriptional programs in Piezo1-deficient bone marrow-derived macrophages [^38^]. Similarly, Luo et al. demonstrated attenuation of liver fibrosis utilizing the same myeloid-specific *Piezo1* knockout [^39^]. Together, these observations implicate mechanotransductive pathways as critical mediators of macrophage responses to fibrotic stiffening, and their sustained expression in DMM suggests a potential feedback loop that reinforces chronic inflammation and ECM remodeling.

Together, our findings provide mechanistic insight into how evolving microenvironmental mechanics may drive synovial pathology. We show that tissue stiffening drives mechano-activation of synovial fibroblasts and macrophages triggering fibrotic transcriptional programs and maladaptive immune-fibroblast crosstalk that potentiates disease. This work broadens our understanding of synovial pathobiology in OA but also identifies cellular and molecular targets for therapeutic intervention aimed at restoring synovial homeostasis.

## Acknowledgements

We would like to thank the research staff at the CHOP CAG Sequencing Core and the Penn Center for Musculoskeletal Disorders (NIH/NIAMS P30AR069619) for their valuable contributions.

## Data Sharing Statement

Datasets used for the present study are available from the corresponding author upon reasonable request.

## Supplementary Materials

### SUPPLEMENTARY METHODS

#### scRNA-seq Data Analysis

Read aligned data from CellRanger was further processed using Seurat (R, v4.1.0). Cells with less than 350 detected genes and 200 total UMIs (Unique Molecular Identifier) were excluded from further analysis. Cells in which >5% of all genes were mitochondrial-derived (i.e., percent.mt>0.05) were likewise excluded. Next, low-quality cells were further filtered by removing those falling within the bottom 8% for both total UMI counts (nCount) and detected gene features (nFeature). After quality control, 13,177 cells were integrated across experimental conditions and *ScaleData()* was used to perform linear transformation and to regress out variation due to cell cycle and mitochondrial gene expression (**Fig 1G**). Unsupervised dimensionality reduction was performed using principal component analysis. Elbow plots and principal component heatmaps were used to determine the number of dimensions for subsequent nonlinear dimensionality reduction via Uniform Manifold Approximation and Projection (UMAP). Clustering was performed using *FindNeighbors* (dims = 1:28) and *FindClusters* (resolution = 0.3) to derive distinct cell clusters. Cluster markers were demarcated using the *FindAllMarkers* function, focusing on genes expressed in greater than 70% of cells in the cluster and in less than 30% of cells in all other cells. For certain cell subpopulations with well-reported markers, clusters markers were identified using violin and gene feature plots.

For higher resolution analysis, fibroblasts were reclustered using *FindNeighbors* (dims = 1:25) and *FindClusters* (resolution = 0.18). Differentially expressed genes (DEGs) between experimental conditions were identified with *FindMarkers* (p_adj_ < 0.05). To functionally annotate fibroblast clusters, DEGs were submitted to PantherDB for statistical overrepresentation analysis using the Biological Process Complete annotation dataset. Pathway enrichment was assessed using Fisher’s Exact test with False Discovery Rate correction, and results were visualized as bubble plots using ggplot2 v3.5.1. *AverageExpression()* was used to generate a pseudobulk expression matrix, which was subsequently inputted for multidimensional scaling, hierarchical clustering and assessment of global changes in transcriptional activity[^40^].

Immune cells were identified through CD45 positivity (i.e., expression of *Ptprc*>1), and those clusters were subsequently subset and reclustered using *FindNeighbors* (dims = 1:20) and *FindClusters* (resolution = 0.37). To evaluate the relative enrichment of pro-inflammatory transcriptional programs, we computed a module score using genes involved in myeloid M1-like (*Il1b, Il6, Tnf, Nos2, Ccl2, Ccl3, Cxcl1, Cxcl2, Cxcl10*) and lymphoid Th1/Th17-like (*Ifng, Il17a, Il23a, Stat1, Tbx21, Rorc*) inflammatory responses [^41–43^]. We likewise calculated pro-fibrotic module scores using genes that have been shown to directly or indirectly induce fibrotic phenotypes phenotypes: *Tgfb1, Il13, Il4, Pdgfb, Pdgfa, Osm, Vegfa, Mmp9, Mmp12, Mmp3, Timp1, Cd163, Csf1r, Mrc1, Cd86, Cd80,* and *Cxcr4* [^44–46^]. These scores were computed via the *AddModuleScore()* function in Seurat.

#### Differential mechanosensitive gene expression

Expression of >30 mechanosensitive genes – including *Piezo-, Trp-, Kcnk-, Tmc-, Asic-, Scnn-, Trek-,* and *Traak*-family genes – was evaluated. Genes with <0.5 average counts in all macrophage clusters were screened out. Remaining genes were compared between DMM and sham macrophages to determine those more prevalent in disease.

#### Trajectory analysis

Macrophages (basal resident, lining, interstitial, and IFNγ-activated) were subset from Seurat and trajectory analysis that was performed in Monocle3 ^[47–49]^. For Sham macrophages (unoperated, sham 4wk, Sham 8wk) and DMM macrophages (DMM 4wk, DMM 8wk), 28 dimensions were included. For clustering, k = 18 and res = 0.001 was applied to sham, while k = 15 and res = 2e-3 was applied to DMM. Default learn_graph parameters were used for both Sham and DMM to identify differentiation trajectories. A wide parameter sweep of learn graph parameters confirmed consistent trends in differentiation. Gene module analysis was performed to identify genes enhanced along pseudotime. Resolutions of 0.019 for sham and 0.01 for DMM were selected, resulting in two modules per condition. Modules with genes enhanced at the end of the pseudotime were submitted for transcription factor motif analysis.

For fibroblasts, sham and unoperated, as well as DMM and unoperated SFs were subset and reclustered in Moncole3. Origins were placed in *Dpp4*+ progenitor SFs present in the unoperated condition. Differentiation trajectories were then computationally determined (sham: Ncenter = 100, minimal_branch_len = 12. DMM: default conditions). Trajectory trends were confirmed to be consistent across multiple parameter combinations.

#### Intercellular stromal-immune crosstalk pathway ranking

The CellChat package (CellChat 1.1.3) [^33^] for R was used to infer and quantify cell-cell communication networks in sham or in DMM synovia at both timepoints. A minimal cell count of ten per cluster was applied. All functions were run with default arguments. Stromal clusters – all fibroblasts, endothelial cells, and pericytes – and immune clusters – macrophages, T cells, B cells, monocytes, and dendritic cells – were defined. Line plots, heatmaps, and river plots were generated for each object with default arguments. A “crosstalk score” was computed for each significant pathway by dividing the probability value of immune-to-stromal or stromal-to-immune interactions by the probability value of all interactions for each pathway. This crosstalk score was then compared between DMM and sham conditions at each time point. Resultant sham proportions were subtracted from corresponding DMM proportions and ranked.

#### Transcription factor motif enrichment

Genes enhanced specifically in the interstitial and IFNγ-activated macrophages were grouped in a module and submitted to RcisTarget [^50^] for transcription factor motif enrichment analysis. The mm9-500bp-upstream-7species.mc9nr.feather file was used for analyses. Significant, high-quality, direct annotation-derived transcription factors (NES > 3, logcount > 2 in >5% of macrophages) that putatively underpinned basal resident macrophage differentiation were identified.

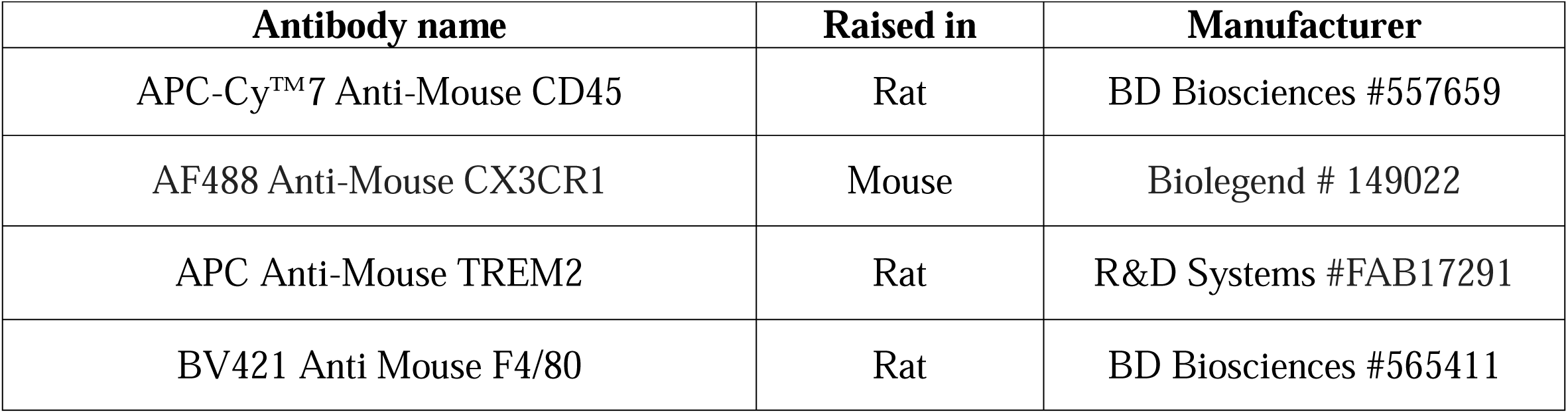
Supplementary Table 1.

## FIGURE LEGEND

**Supplementary Fig 1.** (A) Histopathologic scoring of synovial hyperplasia, cellularity, and fibrosis for anterior femoral [AF], anterior tibial [AT], posterior femoral [PF], and posterior tibial [PT] areas on H&E-stained paraffin sections (n_=_8/group) (B) Feature plots of universal marker genes used to identify synovial fibroblasts. (C) Violin plots showing expression levels of marker genes for each SF subpopulation. (D) Violin and (E) feature plots depicting expression levels of *Thy1* and *Prg4* by distinct SF clusters. (F) Heatmap of expression levels (z-score) of genes involved in the *Mus musculus* GO:2001046 and GO:0032965 pathways. (G) Trajectory analysis of SFs in sham vs DMM synovium (H) Violin plot representing the apoptotic (GO:2001235) to proliferation (GO:0008284) ratio module scores for each SF subcluster (I) Bubble plot showing the expression levels of myofibroblast markers *Col5a3, Lrrc15, Acta2, Tagln* by SF subcluster.

**Supplementary Fig 2.** (A) Representative RNA FISH images localizing lining macrophages, with individual channels displayed independently; dashed lines demarcate the synovial tissue and arrows indicate the resident lining macrophages localized at the intima. (B) Transcript expression of mechanosensitive genes in sham and DMM macrophages

