## Supplementary Materials for "Mechano-activation of synovial fibroblasts and macrophages during OA progression in the dynamically stiffening synovial microenvironment"

**Supplementary Table 1**

| **Antibody name** | **Raised in** | **Manufacturer** |
| --- | --- | --- |
| APC-Cy™7 Anti-Mouse CD45 | Rat | BD Biosciences #557659 |
| AF488 Anti-Mouse CX3CR1 | Mouse | Biolegend # 149022 |
| APC Anti-Mouse TREM2 | Rat | R&D Systems #FAB17291 |
| BV421 Anti Mouse F4/80 | Rat | BD Biosciences #565411 |

**FIGURE LEGEND**

**Fig 1 Altered microenvironmental mechanics and cellular landscape in OA synovium**. (A) Schematic of experimental design. (B) Representative hematoxylin and eosin (H&E) images of the anterior femoral synovial membrane. (C) Atomic force microscopy (AFM) nanomechanical mapping on cryo-sections of murine knee synovium (*n*= 10 mice per group, 157-202 total indentations per group). (D) Representative indentation force versus depth (*F-D*) curves obtained on Sham and DMM synovia 4 weeks post-surgery (measured in PBS with protease inhibitors, 10µm/s rate). Red and blue solid lines depict finite thickness-corrected Hertz model fitted to the entire loading portion (dotted line) of the DMM and Sham *F-D* curves, respectively. Dashed lines represent the unloading portion of *F-D* curves. (E) Superplot of the micromoduli of the synovial membrane (n=10/condition) (F) Fold change in average synovial micromoduli between Sham and DMM joints for each timepoint (G) UMAP projection of all synovial cells from Sham-, DMM-, and un-operated knees (13,177 cells, n=15 biological replicates per condition) (H) Proportional breakdown and (I) total abundance of major synovial cell types in each condition.
